# LC-UV/RI-MS^2^ as the analytical platform for bioconversion of sustainable carbon sources: a showcase of 1,4-butanediol plastic monomer degradation using *Ustilago trichophora*

**DOI:** 10.1101/2023.08.16.553358

**Authors:** An N.T. Phan, Lisa Prigolovkin, Lars M. Blank

**Affiliations:** Institute of Applied Microbiology - iAMB, Aachen Biology and Biotechnology – ABBt, RWTH Aachen University, Worringerweg 1, 52074 Aachen, Germany

## Abstract

Plastic usage by microbes as a carbon source is a promising strategy to increase the recycling quota. 1,4-butanediol (BDO) is a common monomer derived from polyesters and polyurethanes. It presents in the complex mixture from the plastic degradation process. In this study, *Ustilago trichophora* was found to be an efficient cell-factory to valorize BDO. To investigate product formation by *U. trichophora*, we refined the traditional ion exclusion liquid chromatography method by examining eluent, eluent concentrations, oven temperatures, and organic modifiers to make the chromatography compatible with mass spectrometry. An LC-UV/RI-MS^2^ method is presented here to identify and quantify extracellular metabolites in the cell cultures. With this method, we successfully identified that *U. trichophora* secreted malic acid, succinic acid, erythritol, and mannitol into the culture medium. Adaptive laboratory evolution followed by medium optimization significantly improved *U. trichophora* growth on BDO and especially malic acid production. Overall, the carbon yield on the BDO substrate was approximately 33% malic acid. This is the first report on a Ustilaginaceae fungus that was able to convert BDO into versatile chemical building blocks. Since *U. trichophora* is not genetically engineered, it is a promising microbial host to produce malic acid from BDO, thereby contributing to the development of the envisaged sustainable bioeconomy.

## Introduction

Polyurethanes (PUs) are ranked as the 6^th^ most-produced synthetic polymer family. They are synthesized by the reaction between polyols and polyisocyanates. Due to the variability in monomer combinations, the physical properties of PUs can be tailored for a broad range of applications [1]. The PUs market size was valued at 78 billion USD in 2023, which is expected to reach approximately 105 billion USD in 2030 [2]. Consequently, a large amount of PU waste is also being generated and requires proper disposal. Many studies showed that it is feasible to use microorganisms for the biodegradation of PUs, yet at a very low rate [3]. In general, the plastic biodegradation processes focus on degrading plastic polymer chains into oligo- and monomeric compounds, which will be recycled and/or upcycled into value-added products using biotechnology [4]. In the case of PUs, degradation products consist of a broad spectrum of chemicals including amines, alcohols, acids, aromatics, and other compounds (*i*.*e*., ethylene glycol, BDO, adipic acid, 4,4′-ethylenediamine, or 2,4′-toluene diamine), which makes it challenging for post-processing. Therefore, efficient valorization approaches for PU degradation products are in urgent demand to improve the PUs end-of-life processing as well as to achieve a long-term sustainable bioeconomy [4-7].

Plastic biodegradation using microorganisms is still an emerging field. A common challenge lies in the unpredictable behavior of an organism when the carbon source is changed. Furthermore, a new microorganism is discovered to convert a substrate of interest, but no information on its metabolism is available. To overcome these challenges, one option is to use analytical platforms to monitor substrate consumption and identify as well as quantify product formation. Ion exclusion liquid chromatography (IELC) coupled with ultraviolet/refractive index (UV/RI) detectors has been the preferred method in the field of applied microbiology for separating neutral and weakly ionized molecules such as sugars, organic acids, and alcohols [8, 9]. However, the biggest drawback of this method is the difficulty in recognizing overlapping peaks or the ability to identify unknown substances. Furthermore, the fact that sulfuric acid is commonly added to the eluent makes it even harder to adapt this method to any sensitive equipment for peak identification [10]. On the other hand, mass spectrometry (MS) is optimal for peak identification, but it has a limitation on dynamic range and the choices of the mobile phase. Here, we aimed to refine the traditional IELC method to make it compatible with MS. After finding a suitable eluent, we demonstrated the effect of eluent concentrations, oven temperatures, and organic modifiers on peak separations. Our findings allow researchers to customize the method according to their studies. Afterward, an LC-UV/RI-MS^2^ method, which generated comparable peaks from UV/RI and MS, was established to identify and quantify cellular metabolites. Notably, a targeted multiple reaction monitoring (MRM) method covering 40 common compounds in the field of applied microbiology was developed.

In this study, we investigated microorganisms to convert 1,4-butanediol (BDO), one of the monomers from the depolymerization process of polyurethane or polyesters (*i*.*e*., Polybutylene succinate or Polybutylene adipate terephthalate) [11]. BDO is quite a challenging substrate because it is not naturally produced by any known microorganism. Hence, the fungal family Ustilaginaceae has been drawing a lot of attention due to its capability of using a wide range of sustainable substrates to produce valuable chemicals of industrial interest [12]. Therefore, Ustilaginaceae could be potential candidates for BDO degradation. From a screening among Ustilaginaceae species, we found that *U. trichophora* was able to degrade BDO naturally. Adaptive laboratory evolution (ALE) has been performed to improve cell growth. By using our newly developed LC-UV/RI-MS^2^ method, malic acid, succinic acid, mannitol, and erythritol were identified as the dominant extracellular metabolites. Finally, we successfully improved malic acid production, titer, and substrate-to-product yield by optimizing medium compositions. When we supplemented the medium with CaCO_3_, the product precipitated in the form of calcium malate, and facile isolation from the cell culture is achieved. This is a tremendous advantage in downstream processing when we utilize waste as substrate.

Due to the economic benefit of BDO in plastic production, previous research mainly focused on its production. Our findings will contribute toward the conception of bio-upcycling plastic waste by connecting the biodegradation of mixed plastic waste to the selective biosynthesis of valuable chemicals in microbial cell factories. Moreover, the use of non-genetically modified organism can broaden the applications as we can avoid consumer hesitance and regulatory restrictions.

## Experimental Procedures

### Method optimization for IELC-HPLC

Two independent HPLC systems were used to avoid cross-contamination. A Dionex UltiMate 3000 HPLC System (*Thermo Scientific*, Germany) was applied for the mobile phases containing H_2_SO_4_ (5 mM) and TFA (0.1%). Signals were detected with a Dionex UltiMate 3000 Variable Wavelength Detector and a SHODEX RI-101 detector (*Showa Denko Europe GmbH*, Germany). When using formic acid (FA) as an additive, experiments were performed with a Shimadzu Nexera UHPLC system coupled with a RID-20A Refractive Index detector and an SPD-40 UV detector (*Shimadzu*, Japan). The cell temperature of all UV and RI detectors was set to 40 °C. The UV detectors were set at 210 nm. Two identical columns Isera Metab-AAC 300 × 7.8 mm column (*Isera*, Germany) were installed in both HPLC. The injection volume was 5 μL. The flow rate was 0.4 mL/min, and the column oven was set at 40 °C unless otherwise noted. The whole system was purged and equilibrated when a parameter was changed before analysis. An authentic standard mixture was used in all optimization steps. This mixture included 20 g/L cellobiose, 20 g/L glucose, 20 g/L quinic acid, 20 g/L xylitol, 20 g/L glycolaldehyde, 20 g/L itaconic acid, 10 g/L adipic acid, 20% MeOH, and 20 g/L BDO. All chemicals were obtained from Carl Roth GmbH & Co. KG (Germany) or Sigma Aldrich Chemie GmbH (Germany) with a purity higher than 99%.

### LC-UV/RI-MS^2^

5 μL of each sample were injected into an Isera Metab-AAC 300 × 7.8 mm column (*Isera*, Germany) with a Shimadzu Nexera UHPLC (*Shimadzu*, Japan). Unless otherwise noted, experiments were performed with an FA concentration of 0.2%, a flow rate of 0.4 mL/min, a column temperature of 40 °C, and without organic modifiers. After the samples passed the column, the flow was split into two directions with a split ratio of 1 to 10. A major part of the samples was measured with a RID-20A Refractive Index detector and an SPD-40 UV detector at 210 nm (*Shimadzu*, Japan). The cell temperatures of UV and RI detectors were 40 °C. The rest was analyzed with a triple quadrupole MS 8060 (*Shimadzu*, Japan). The mass spectrometric parameters were: ESI negative and positive mode; scan speed of 15,000 u/sec; desolvation line temperature: 250 °C; nebulizer gas flow: 3 L/min; heat block temperature: 400 °C; other parameters were optimized automatically by auto-tuning. An MRM transition library of 40 metabolites was constructed using authentic standards (Table S3). Authentic standard mixtures were injected periodically throughout the analysis to evaluate the stability of the analytical system and to support peak identification.

For quantification (Table S2 and S3), each metabolite was analyzed at 16 different concentrations with a dilution factor of 2 at each step. All compounds were prepared with concentrations ranging from 0.001 -20 g/L, except for the concentrations of fumaric acid, itaconic acid, and aconitic acid were 0.0003 -5 g/L. Data were processed with LabSolutions software version 5.97 (*Shimadzu*, Japan). All chemicals were obtained from Carl Roth GmbH & Co. KG (Germany) or Sigma Aldrich Chemie GmbH (Germany) with a purity higher than 99%.

### Strain and culture conditions

Modified Tabuchi Medium (MTM) was used according to Geiser *et al*. with 0.2 g/L MgSO_4_·7H_2_O, 10 mg/L FeSO_4_·7H_2_O, 0.5 g/L KH_2_PO_4_, 0.8 g/L NH_4_Cl, 1 mL/L vitamin solution and 1 mL/L trace element solution [12]. The vitamin solution contained 0.05 g/L d-biotin, 1 g/L d-calcium pantothenate, 1 g/L nicotinic acid, 25 g/L myo-inositol, 1 g/L thiamine hydrochloride, 1 g/L pyridoxol hydrochloride, and 0.2 g/L *para*-aminobenzoic acid. The trace element solution contained 1.5 g/L EDTA, 0.45 g/L of ZnSO_4_·7H_2_O, 0.10 g/L of MnCl_2_·4H_2_O, 0.03 g/L of CoCl_2_·6H_2_O, 0.03 g/L of CuSO_4_·5H_2_O, 0.04 g/L of Na_2_MoO_4_·2H_2_O, 0.45 g/L of CaCl_2_·2H_2_O, 0.3 g/L of FeSO_4_·7H_2_O, 0.10 g/L of H_3_BO_3_, and 0.01 g/L of KI. A two-step preculture was performed with 50 g/L glucose and then with the indicated BDO concentrations. 100 mM of 2-(*N*-morpholino)ethanesulfonic acid (MES) or 33 g/L of CaCO_3_ were utilized as buffers. All medium components were obtained from Carl Roth GmbH & Co. KG (Germany) or Sigma Aldrich Chemie GmbH (Germany).

*U. maydis* MB215 [13], *U. trichophora* TZ1 [14], *U. cynodontis fuz7 [15]* and *U. vetiveriae* TZ1 [16] were used in this study. For adaptive laboratory evolution (ALE), *U. trichophora* TZ1 was grown in MES-MTM with 40 g/L BDO in 100 mL Erlenmeyer flasks with 10% (v/v) filling volume. OD_600_ was measured every 2-3 days, after which a new culture was inoculated to an OD_600_ of 0.5. This procedure was repeated sequentially for 74 days.

Cell growth was screened using the Growth Profiler GP960 (*EnzyScreen*, The Netherlands). Strains were cultivated in grey polystyrene square 24-deep-well microplates with a transparent bottom and a filling volume of 1.5 mL per well (225 rpm, d = 50 mm). Time-course pictures were taken and processed using EnzyScreen Image software to generate green values, which are correlated with cell biomass OD_600_ (Figure S1).

All other cultivations were performed in System Duetz^®^ 24-deep-well plates (*EnzyScreen*, The Netherlands) with 1.5 mL MTM per well, incubated at 30 °C, relative air humidity of 80%, and a shaking speed of 300 rpm (*Infors HT Multitron Pro* shaker) [17]. The cultures were prepared in parallel into multiple plates and each plate was sacrificed for a data point to ensure constant oxygenation. This experimental design has been widely applied for microorganism cultivation including the fungus in the Ustilaginaceae family as well as *U. trichophora* [18-20].

For HPLC analysis, all samples were filtered with Rotilabo^®^ (CA, 0.2 μm, Ø 15 mm) or Acrodisc^®^ Syringe Filters (GHP, 0.2 μm, Ø 13 mm) and diluted in total 2-fold with water. In case of using CaCO_3_ as a buffer, samples were dissolved 1:1 with 2 M HCl.

## Data processing

Heatmap was conducted using Multiexperiment View Version 4.9 [21] (Dana-Farber Cancer Institute, USA, available at http://www.tm4.org/mev).

## Results and Discussion

### LC-UV/RI-MS^2^ method for extracellular metabolite analysis

The widely employed IELC eluents are usually diluted mineral acids or strong organic acids, which are incompatible with MS due to their low volatility and the risk of damaging the MS interface [10]. Volatile eluents like acetic acid, formic acid (FA), or trifluoroacetic acid (TFA) have been tested to replace sulfuric acid; however, the applications were mostly limited to the separation of organic acids [22-25]. Therefore, the first step of refining IELC was to identify a suitable acidic mobile phase. Here, we tested the separation of a standard mixture in three different conditions including 5 mM H_2_SO_4_, 0.1% TFA, and 1% FA. All eluents had a pH of approximately 2.1 at 22 °C. The representative standard mixture consisted of a disaccharide (cellobiose), a monosaccharide (glucose), a cyclohexane-carboxylic acid (quinic acid), a sugar alcohol (xylitol), an alcohol aldehyde (glycolaldehyde), an unsaturated dicarboxylic acid (itaconic acid), a dicarboxylic acid (adipic acid), an alcohol (MeOH), and a diol (BDO). We observed that all metabolites in the standard mixture were separated quite well in all eluents (Figure 1). With the final goal to develop an LC-UV/RI-MS^2^ method, the mobile phase should be compatible with all detectors. Considering the MS, FA was the better choice, because TFA is well known for causing ion suppression at levels exceeding 0.01% [26]. Another important point that should be considered is the acid concentration. Carboxylic acids have the same cut-off wavelength at 210 nm. The higher the organic acid concentration in the mobile phase, the less sensitive the signal is in UV/RI detectors. With the current FA concentration of 1%, no organic acids can be observed in the UV chromatogram except for itaconic acid (Figure S2). Therefore, the HPLC method needed to be further optimized.

**Figure 1.**
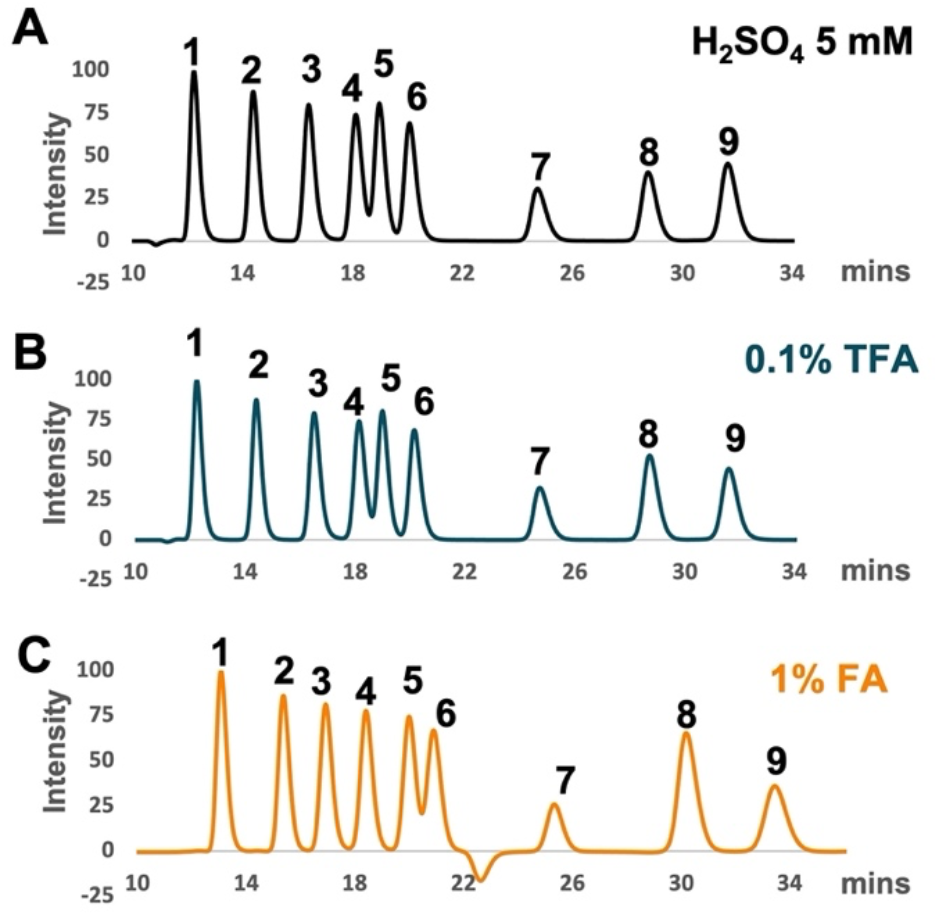
Ion exclusion chromatography with different mobile phases: (A) 5 mM H_2_SO_4_; (B) 0.1% TFA; and (C) 1% FA. The standard mixtures contained (1) cellobiose; (2) glucose; (3) quinic acid; (4) xylitol; glycolaldehyde; (6) itaconic acid; (7) adipic acid; (8) MeOH, and (9) BDO. Peak intensities were normalized to the highest peak in each condition.

First, experiments were performed with the FA concentration in eluents from 1% to 0.2%. The priority was to decrease the FA concentration as much as possible to improve the sensitivity of UV signals but to keep accurate peak separation. We observed scarcely changes in peak retention times (Figure 2A), whereas the background signals dropped and the sensitivity from the UV detector was significantly improved (Figure S2). Among all tested metabolites, organic acids were influenced the most by the eluent acidity (Figure 2B). This result was expected because variations in the eluent pH could alter the degree of ionization of the metabolites and therefore alter the retention times [27]. Notably, the respective organic acids responded differently to the varied acid concentration. While the retention of quinic acid decreased, itaconic acid and adipic acid retention times increased, respectively, with the FA reduction in the mobile phase.

**Figure 2.**
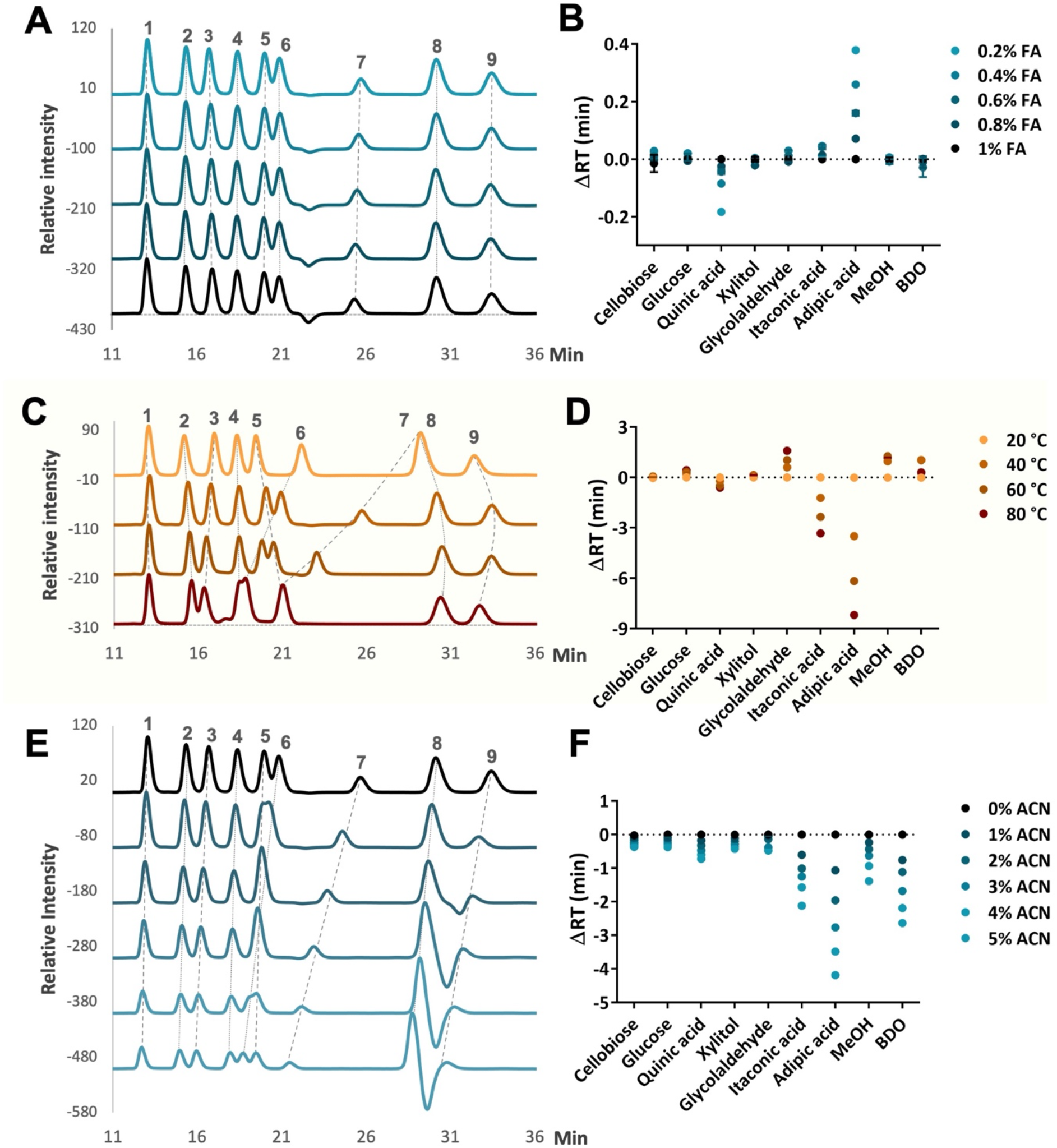
Effects of HPLC parameters on metabolite retention times. (A, B) FA concentration, (C, D) temperature, and (E, F) ACN concentration. (A, C, and E) RI chromatograms with peak intensities normalized to the highest peak in each run. (B, D, and F) The changes in retention time compared to reference conditions included 1% FA, 20 °C, and 0% ACN, respectively. Each data point represents the average of 3-4 technical replications, with the error bar indicating the standard deviation. The standard mixtures contained (1) cellobiose; (2) glucose; (3) quinic acid; (4) xylitol; (5) glycolaldehyde; (6) itaconic acid; (7) adipic acid; (8) MeOH, and (9) BDO. Except when a specific condition was tested, experiments were performed with an FA concentration of 0.2%, a flow rate of 0.4 mL/min, a column temperature of 40 °C, and no organic modifiers.

Like other HPLC methods, raising column temperature is a facile way to influence peak separations by reducing analyte retentions and improving peak shapes [27, 28]. Moreover, the high temperature was suggested to elevate the ionization level in IELC [27]. A resin-based column is advantageous in a wide range of working temperatures. Therefore, the standard mixture was tested at four different temperatures ranging from 20 °C to 80 °C (Figure 2C, D). The increased temperature had a negative impact on all organic acid retentions.

This result was in agreement with previous studies on several aliphatic and aromatic acids [29]. Other metabolites showed a different behavior. While cellobiose and xylitol exhibited modest changes, glucose, and glycolaldehyde were retained longer at elevated temperatures. We observed that the correlation between the temperature variations and the retentions of MeOH and BDO was not linear. This might be because of hydrogen bonding between the non-ionic molecules and the sulfonate groups on the stationary phase [30]. Unlike organic acids, other metabolites in the standard mixture were either neutral or weakly ionized molecules. When the column was heated, the viscosity of the mobile phase and samples decreased. Thus, they were able to penetrate deeper into the resin interior and retained longer on the column.

Though the major eluents used in IELC was water, several studies found that the addition of organic modifiers can change the metabolite dissociation constants and affect retention by alteration of the chromatographic efficiency [27, 30]. The column stationary phase is polystyrene-divinylbenzene (PS/DVB) with 8% H^+^ crossed-linked, which is very sensitive to organic solvents. From the manufacturer’s recommendation, this type of column should not be operated with more than 30% of ACN or 5% of alcohol (*i*.*e*., MeOH or isopropanol). Because alcohols might shrink the resin and cause irreversible damage to the column [31], we tested the mobile phase with the addition of ACN ranging from 0% to 5%. Though the degree of effects on each compound was different, we observed the retentions of all metabolites decreased with the increase in ACN level (Figure 2E, F). In case proper separation has already been achieved, an organic modifier is helpful to reduce the analysis time.

To validate this method, the standard mixture was diluted two times and analyzed by HPLC-UV/RI with an FA concentration of 0.2%, a flow rate of 0.4 mL/min, a column temperature of 40 °C, and without organic modifiers (supplemental experiment procedures; table S1). The relative standard deviation (RSD) values of intra-day samples were between 0.3% and 2.5%, with the accuracy ranging from 95% to 100% for all analytes. For inter-day samples, the RSD values showed between 5% and 10%, with accuracy varying between 98% and 104%. All results were below the ±15% threshold [32], which indicated that our method is robust and accurate for future applications.

Ultimately, we designed an LC-UV/RI-MS^2^ method to support the identification of unknown or overlapped peaks in the fermentation broth and cultivation media. Once the samples were separated on the column, the flow was split into two directions with a split ratio of 1 to 10. A major part of the sample was measured with UV/RI and the rest was analyzed with MS. With this configuration, we generated comparable peaks from UV/RI and MS. The IELC was performed with an FA concentration of 0.2%, a flow rate of 0.4 mL/min, a column temperature of 40 °C, and without organic modifiers.

As the first attempt to support peak identification, an MRM library was developed for 40 organic compounds common in applied microbiology as either substrates or by-/side products (Table S3). All conditions were chosen to acquire proper peak identification regarding the combination of chromatographic retention time and accurate mass differences.

Altogether, there was no generally applicable rule that increasing or decreasing a parameter could improve peak retention or resolution. The effects of a parameter on a given analysis depended not only on the individual characteristics of an analyte but also on the composition of different metabolites in the mixture. The results from this study provide a general overview of IELC conditions including eluent pH, temperatures, and the organic modifier composition, which can be adjusted either individually or in combination to enhance resolution and selectivity. By not changing the resin stationary phase compared to the traditional method, this LC-UV/RI-MS^2^ method can be easily adapted in any laboratory currently using the traditional IELC method with UV/RI detectors. Moreover, the use of a triple quadrupole MS is advantageous in being affordable, efficient, and easy to operate compared to other high-resolution platforms [33]. The eluent is aqueous acid with no organic modifier, which offers environmental and economic benefits in addition to compatibility with aqueous sample matrices.

### Selection and evolution of Ustilaginaceae for bioconversion of BDO

Ustilaginaceae have been drawing a lot of attention in biotechnology due to their ability to use many sustainable substrates as feedstock and produce valuable chemicals of industrial interest [12]. As the bioconversion of BDO is not well understood, Ustilaginaceae fungi became potential candidates to investigate whether they had the ability to use BDO as the sole carbon source. *U. maydis* MB215 [13], *U. trichophora* TZ1 [14], *U. cynodontis* Δ*fuz [15]*, and *U. vetiveriae* TZ1 [16] were chosen because they naturally produced many value-added products and their genomes were sequenced. Experiments were performed using MTM minimum medium with 30 g/L BDO. Interestingly, among four Ustilaginaceae strains, only *U. trichophora* TZ1 was able to grow on BDO (Figure 3A). This result indicated that *U. trichophora* TZ1 had the fundamental metabolic pathways to degrade BDO compared to other Ustilaginaceae strains.

**Figure 3.**
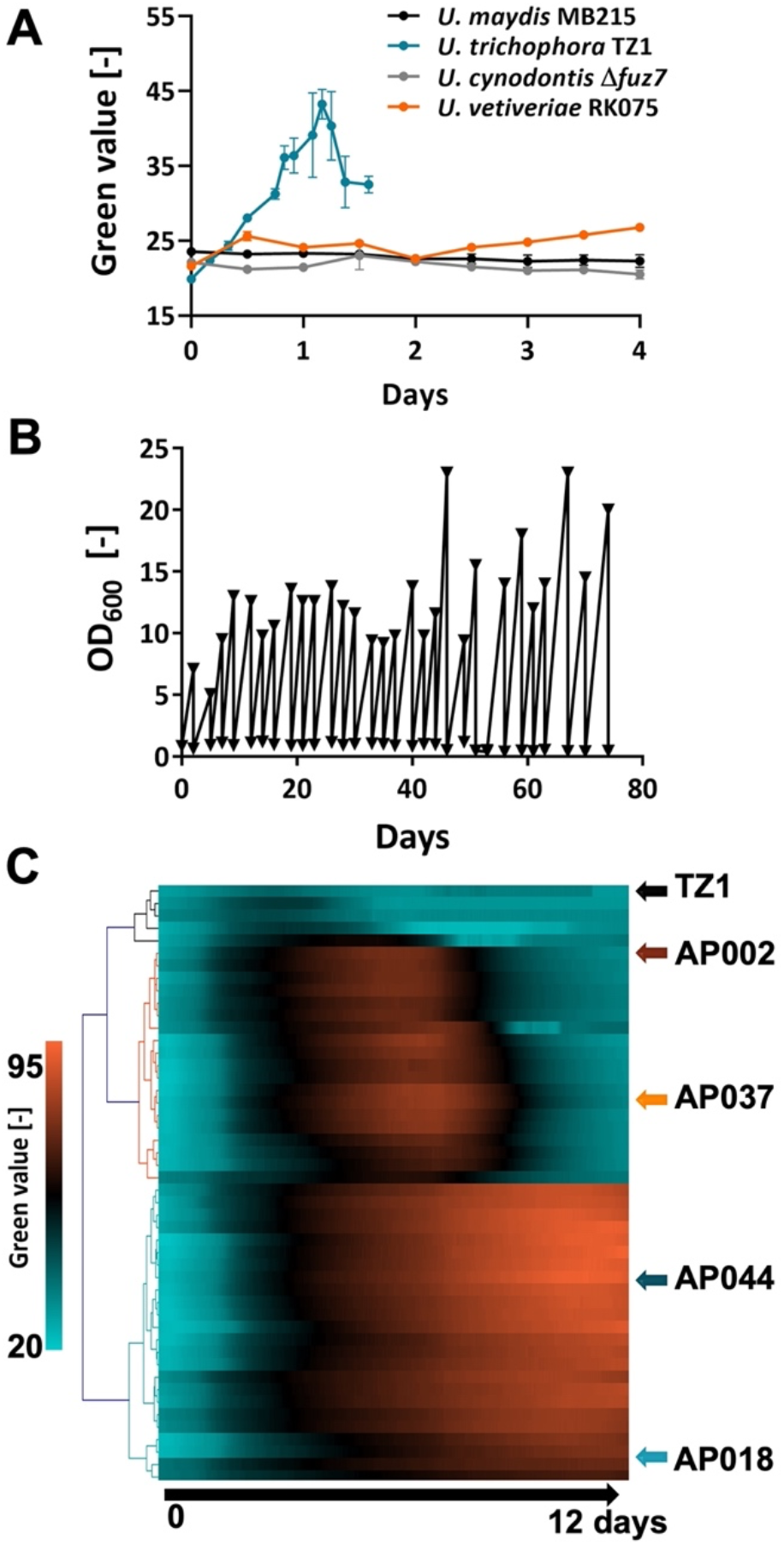
Screening Ustilaginaceae strains for BDO consumption. (A) The growth of *U. maydis* MB215, *U. trichophora* TZ1, *U. cynodontis* Δ?*fuz7, and U. vetiveriae* TZ1 on MTM with BDO as sole carbon source. Experiments were performed in duplicates. The error bars indicate the standard deviation. (B) ALE of *U. trichophora* TZ1 on MTM-BDO. (C) The heatmap for growth of the control strain *U. trichophora* TZ1 and 46 random strains on MTM-BDO after the ALE experiment (n=1).

To improve the growth of *U. trichophora* TZ1, we performed adaptive laboratory evolution (ALE) on this strain with MTM-MES medium containing 40 g/L BDO (Figure 3B). This method has been proven as a powerful approach using evolutionary forces to influence strain phenotypes, performance, and stability [34]. After 74 days with 28 re-inoculations, cell growths were significantly improved. The growth of the control strain *U. trichophora* TZ1 and 46 random strains from evolved cultures were shown in the heatmap (Figure 3C). We observed three clusters representing different growing trends. Four strains did not show any improvement in growth and were grouped with the control strain (upper cluster). The other strains presented significantly improved growth compared to *U. trichophora* TZ1. While the strains in the bottom cluster exhibited the rise of green values over time, the green values of strains in the middle cluster increased up to 6-8 days and then decreased.

To our knowledge, this is the first report on a fungus that uses BDO as the sole carbon source. When *U. trichophora* TZ1 used glycerol as a carbon source, the green value correlated with OD_600_ (Figure S1). However, in this experiment, we observed that the color of the medium changed to brown for the strains in the middle cluster. Therefore, we could not directly convert the green values into OD_600._ Two represented strains from each middle and lower cluster, AP002, AP037, AP044, and AP018 were selected for further investigation (Figure 3C).

### Identification and quantification of extracellular metabolites

To gain better knowledge of the differences among evolved strains, further investigations were required with biological replicates. Evolved strains were cultivated with MTM-MES medium containing 40 g/L BDO (Figure 4). Time-course sampling was performed to measure cell density and quantify extracellular metabolites. First, we aimed to identify which metabolites were produced extracellularly while *U. trichophora* consumed BDO. The cell cultures after four days were analyzed with the new LC-UV/RI-MS^2^ method. Besides the substrate BDO, three unknown peaks were detected in RI chromatograms (Figure 4A). With our MRM method, the first and second peaks were identified as malic acid and mannitol, respectively. Notably, we found that succinic acid and erythritol coeluted at a retention time of 19 minutes. The identifications of all metabolites were confirmed with authentic standards. All four peaks were separated by increasing the column temperature from 40 °C to 75 °C (Figure 4B). This improvement allowed the use of only UV/RI detectors for quantification in further steps. In general, those results indicated that the new LC-UV/RI-MS^2^ method is powerful for identifying unknown peaks and recognized coeluted peaks, which would be challenging if only the conventional UV/RI method was applied.

**Figure 4.**
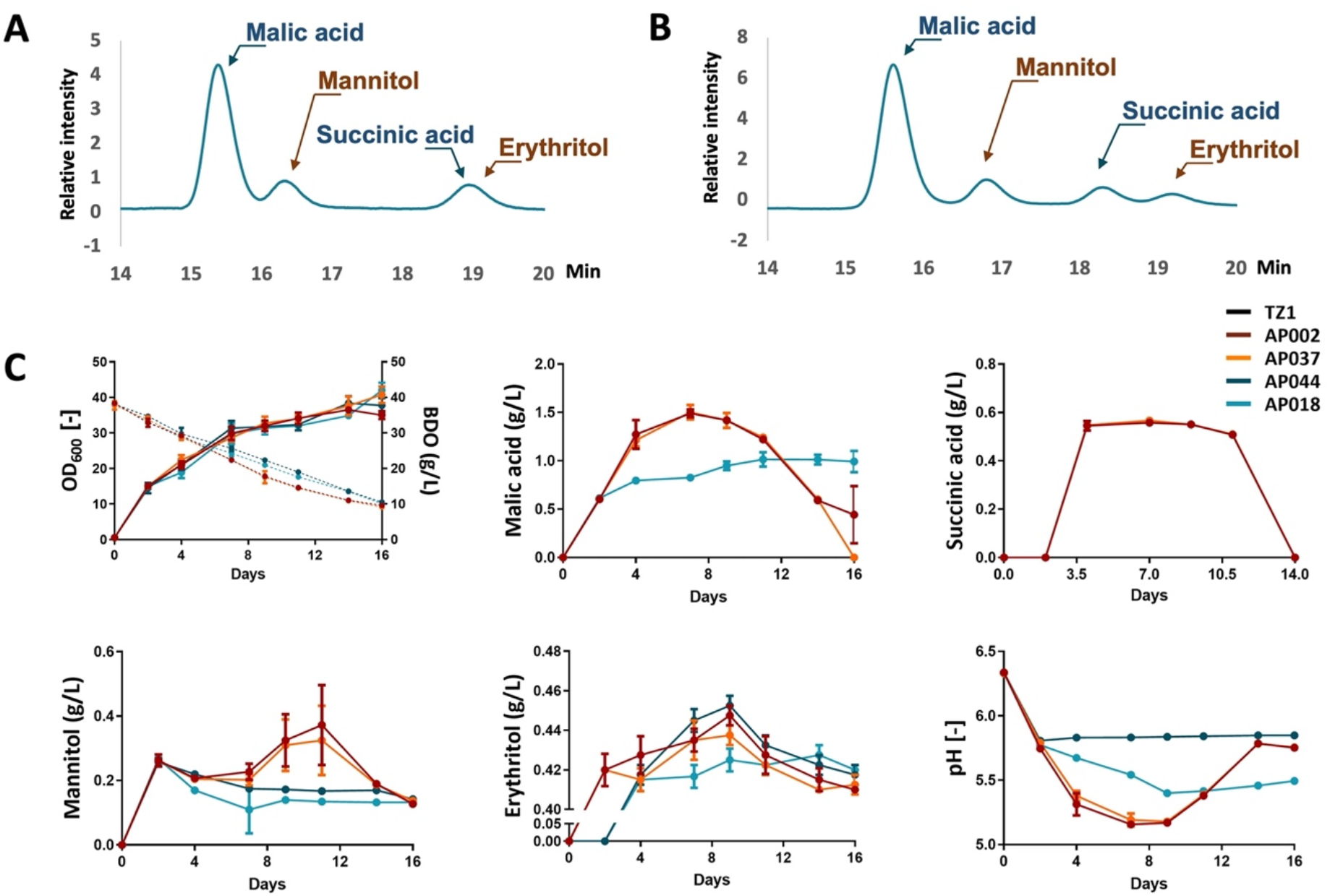
*U. trichophora* is the novel candidate for bioconversion of BDO. Four evolved strains namely AP002, AP037, AP044, and AP018 were cultivated with MTM-MES medium containing 0.4 M BDO. (A) LC-UV/RI-MS^2^ was used to identify the extracellular metabolites with an FA concentration of 0.2%, a flow rate of 0.4 mL/min, a column temperature of 40 °C, and without organic modifiers. (B) Peak separation was improved by increasing the oven temperature from 40 °C to 75 °C. The representative RI chromatograms were from the culture of AP002 after 4 days of cultivation. (C) Time course sampling was performed to measure OD_600_ and pH values as well as the concentrations of BDO, malic acid, succinic acid, mannitol, and erythritol. The error bars indicate the standard deviation. Experiments were performed with four biological replicates for each strain.

All four evolved strains showed similar growths but significantly different extracellular metabolic profiles (Figure 5C). Interestingly, AP002 and AP037 produced higher levels of organic acids than other strains. The highest titer of malic acid and succinic acid from AP037 was 1.50 ± 0.08 g/L and 0.57 ± 0.01 g/L, respectively. In addition, all strains produced mannitol and erythritol. The highest abundance of mannitol was 0.37 ± 0.12 g/L from AP002 and erythritol was 0.45 ± 0.01 g/L from AP044.

**Figure 5.**
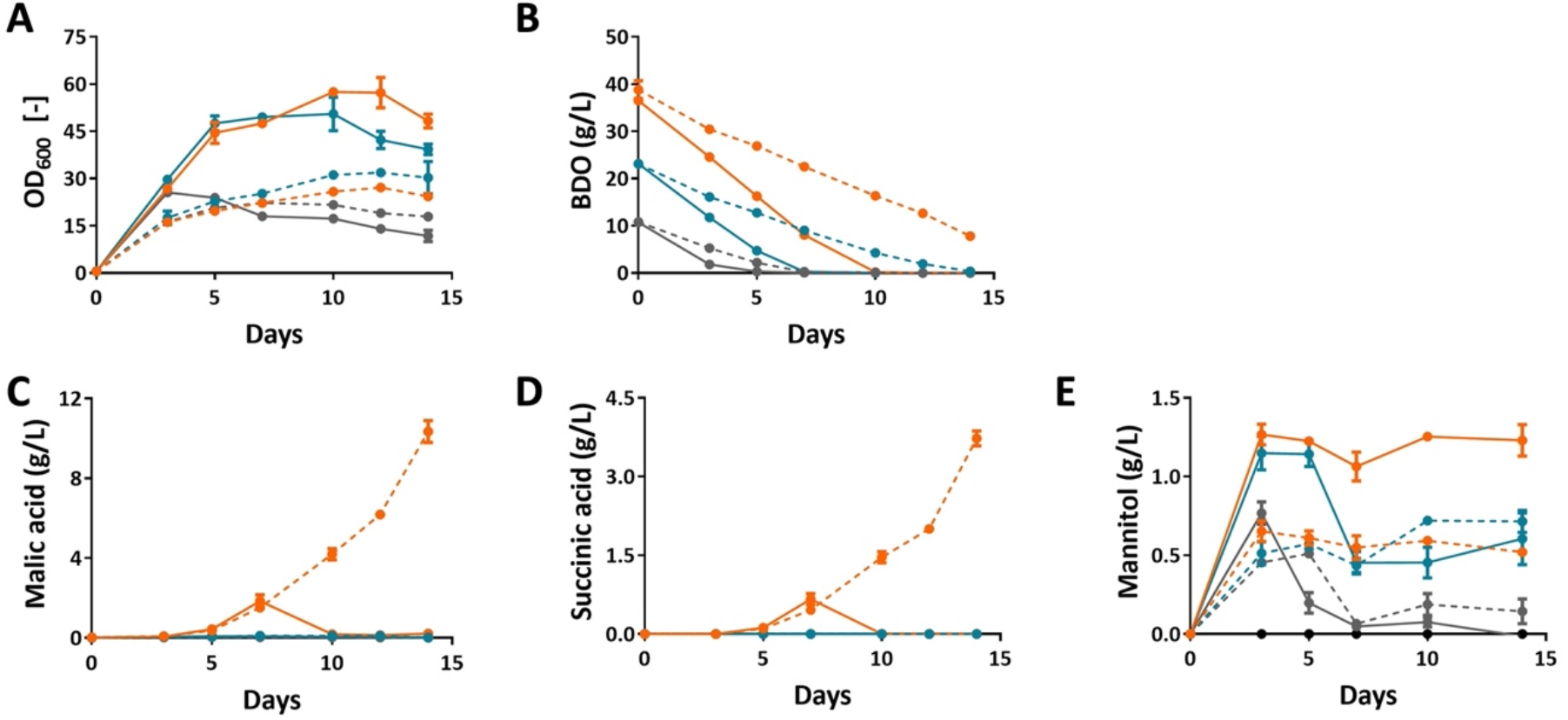
Medium optimization to improve malic acid production from BDO. Strain AP037 was grown on double MTM medium (solid line) and single MTM medium (dashed line). In double MTM medium, the concentrations of all components were double except for BDO and CaCO_3_. Different BDO concentrations were tested including 10 g/L (grey), 25 g/L (blue), and 40 g/L (orange). In all conditions, 33 g/L CaCO_3_ was used as a buffer. Time course sampling was performed to measure (A) OD_600_, (B) BDO consumption, as well as (C) malic acid, (D) succinic acid, and (E) mannitol production. The error bars indicate the standard deviation. Experiments were performed with four biological replicates for each condition.

In summary, we found that *U. trichophora* was not only capable of degrading BDO but also of producing add-value compounds including malic acid, succinic acid, mannitol, and erythritol. From previous studies, *U. trichophora* is well-known for producing malic acid and succinic acid from glycerol [14]. This is the first report confirming mannitol and erythritol production in *U. trichophora*.

### Medium optimization increases malic acid production from BDO

Malic acid, one of the top value-added building block compounds from biomass [35], was the most abundant extracellular metabolite when *U. trichophora* used BDO as the sole carbon source. Therefore, we decided to further improve malic acid production of *U. trichophora* AP037. A previous study showed that medium compositions directly affected malic acid production in *U. trichophora* [16]. Specifically, *U. trichophora* produced more malic acid while cultivating with double MTM medium (doubled concentrations of all medium components except carbon source). Thus, we investigated both standard and double MTM medium. Various initial BDO concentrations were examined including 10 g/L, 25 g/L, and 40 g/L. Because CaCO_3_ has been proven as an effective additive in organic acid production, we decided to test CaCO_3_ instead of MES in all experiments.

While comparing single and double MTM medium, we observed clear differences in biomass formation. *U. trichophora* consumed BDO faster and cell growth rates were almost two times higher in double MTM medium (Figure 5A, B), most likely due to a higher level of the nitrogen source. This result was supported by a previous study that showed that *U. trichophora* only produced malic acid after nitrogen depletion [14, 18, 19]. For example, under the same initial BDO concentration of 10 g/L, the growth rates were 0.22 ± 0.00 h^-1^ and 0.35 ± 0.01 h^-1^ for single and double MTM medium, respectively. Moreover, higher amounts of mannitol were produced in a double MTM medium (Figure 5E). Malic acid and succinic acid were detected only when the initial BDO concentration was 40 g/L (Figure C, D). Hence, with the aim of improving malic acid production, a single MTM medium with 40 g/L BDO was more suitable than a double MTM medium. The highest titer of malic acid was 10.35 ±?0.55 g/L, which was 6.7-fold increase compared to the experiment using MES as the buffer. The total yield was 0.33± 0.03 g_mal_/g_BDO_ (or 0.23± 0.02 mol_mal_/mol_BDO_). This improvement is assigned to the added CaCO_3_, which keeps the pH of the culture medium above 6.5.

In general, we successfully increased the titer and especially the yield of the malic acid production from BDO. When CaCO_3_ was used as an additive, calcium malate precipitated, and the product of interest was isolated out of the culture broth. This feature not only helped to avoid product inhibition but also made it easier for post-processing in the future when the mixture of PUs degradation products is used as a carbon source. With a malic acid production yield of approximately 33%, *U. trichophora* AP037 is a promising microbial host to valorize BDO and broaden the applicability of Ustilaginaceae fungus in plastic upcycling.

## ASSOCIATED CONTENT

Supporting Information

## AUTHOR INFORMATION

Corresponding Author *

## Notes

The authors declare no competing financial interest.

## ACKNOWLEDGMENT

A.N.T.P. has received funding from the European Union’s Horizon 2020 research and innovation programme under the Marie Sklodowska-Curie grant agreement No. 793158. The LC-UV/RI-MS^2^ was funded by the Deutsche Forschungsgemeinschaft (DFG, German Research Foundation) under Germany’s Excellence Strategy within the Cluster of Excellence FSC 2186 ‘The Fuel Science Center’. We would like to thank Shimadzu Europa GmbH for the fruitful collaboration on developing the LC-UV/RI-MS^2^ method.

