## Supplementary material for "LC-UV/RI-MS^2^ as the analytical platform for bioconversion of sustainable carbon sources: a showcase of 1,4-butanediol plastic monomer degradation using *Ustilago trichophora*": Figure S1, Figure S2, Table S1, Table S2, Table S3

#### Supplemental Experimental procedures

##### IELC method validation

Samples were analyzed on three different days in triplicate. The RSD values was calculated by dividing the standard deviation by the measured concentration. The percent accuracy was determined by $[(C_m – C_k)/C_k] * 100\%$, where $C_m$ is the measured concentration and $C_k$ is the known concentration of the analytes in standard mixtures. The method was considered validated if the RSD and accuracy were within 15% of the measured and known concentrations for both the intra-day and the inter-day results.

#### Supplemental figures

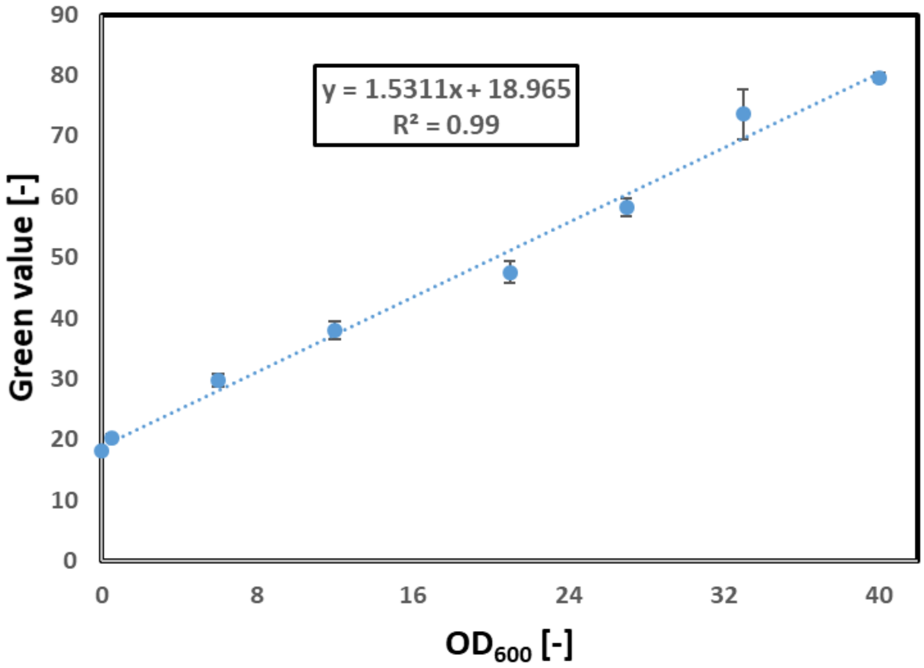


**Figure S1.** Correlation between green value and $OD_{600}$ for *U. trichophora*. *U. trichophora* TZ1 was cultivated with MES-MTM with 50 g/L glycerol as carbon source. Cells were collected after 72h and used to prepare solution with the pre-defined $OD_{600}$ ranging from 0.5 to 40. The sample for $OD_{600}$ of 0 was the medium MES-MTM with 50 g/L glycerol. All solutions with pre-defined $OD_{600}$ were loaded into 24-deep-well microplates and measured the green value by Growth Profiler GP960 (*EnzyScreen*, The Netherlands). Experiments were performed with three biological replicates. The error bars indicate the standard deviation.

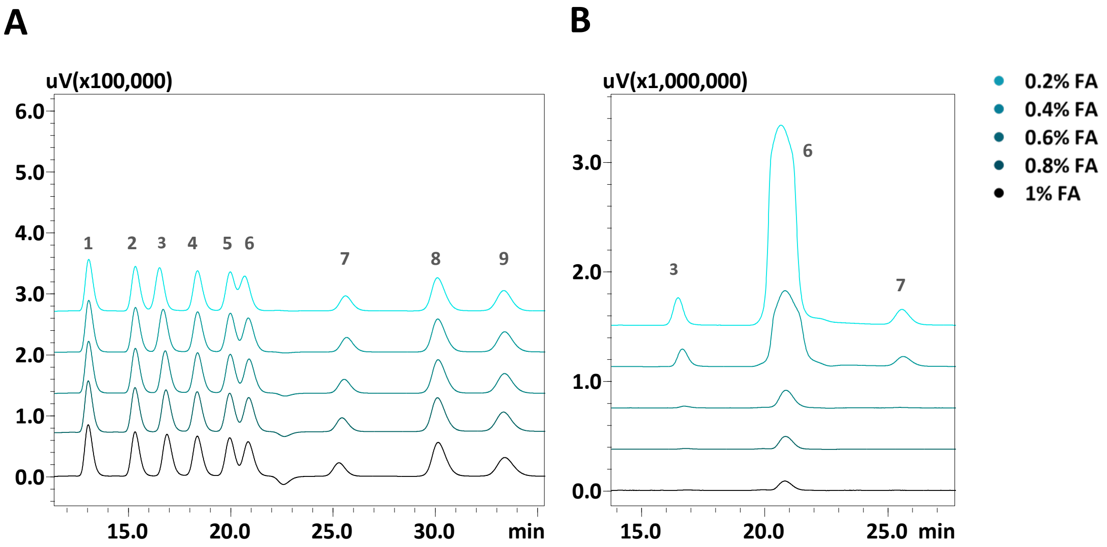


**Figure S2.** Effects of FA concentration. (A) RI chromatograms. (B) UV chromatograms. The standard mixtures contained (1) cellobiose; (2) glucose; (3) quinic acid; (4) xylitol; (5) glycolaldehyde; (6) itaconic acid; (7) adipic acid; (8) MeOH, and (9) BDO. Experiments were performed with a FA concentration from 0.2% to 1%, a flow rate of 0.4 mL/min, a column temperature of 40 °C, and without organic modifiers.

### Supplemental tables

**Table S1:** Method validation

| | $C_k$ (g/L) | Intra-day | | | Inter-day | | |
|---|---|---|---|---|---|---|---|
| | | $C_m$ (g/L) | RSD (%) | Accuracy (%) | $C_m$ (g/L) | RSD (%) | Accuracy (%) |
| **Cellobiose** | 10 | 9,8 | 0.3 | 98 | 10.4 | 8 | 104 |
| **Glucose** | 10 | 9.8 | 0.6 | 98 | 10.3 | 7 | 103 |
| **Quinic acid** | 10 | 9.8 | 0.6 | 98 | 10.3 | 7 | 103 |
| **Xylitol** | 10 | 9.9 | 0.8 | 99 | 10.3 | 6 | 103 |
| **Glycolaldehyde** | 10 | 9.9 | 0.7 | 99 | 10.3 | 5 | 103 |
| **Itaconic acid** | 10 | 9.9 | 1.2 | 99 | 10.3 | 6 | 103 |
| **Adipic acid** | 5 | 4.8 | 2.5 | 95 | 4.9 | 10 | 98 |
| **MeOH** | 10 | 9.5 | 1.8 | 95 | 10.1 | 7 | 101 |
| **BDO** | 10 | 10.0 | 1.3 | 100 | 10.3 | 6 | 103 |

$C_k$: the known concentration of the analytes.
$C_m$: the measured concentration of the analytes.

**Table S2:** UV/RI detectors

| Compound ID | Ret. Time | Quantification | | |
|---|---|---|---|---|
| | | Range (g/L) | Equation | $R^2$ |
| **2-ketogluconic acid** | 12.49 | 0.01 - 20 | y = 323004x + 84916 | 0.99 |
| **Acetic acid** | 23.93 | 0.04 - 20 | y = 318966x + 38214 | 1 |
| **Aconic acid** | 18.80 | 0.08 - 20 | y = 108960x - 2751.5 | 1 |
| **Adipic acid** | 25.36 | 0.01 - 5 | y = 372380x + 27356 | 0.99 |
| **Alpha-ketoglutaric acid** | 12.97 | 0.001 - 5 | y = 3E+06x + 170251 | 0.99 |
| **Arabinose** | 16.94 | 0.08 - 10 | y = 104636x - 18204 | 1 |
| **BDO** | 32.84 | 0.01 - 10 | y = 74267x - 6291.1 | 1 |
| **Cellobiose** | 12.57 | 0.005 - 20 | y = 112227x + 1472.8 | 1 |
| **EG** | 25.03 | 0.01 - 20 | y = 66379x - 2159.4 | 1 |
| **Erythritol** | 18.61 | 0.02 - 20 | y = 101941x + 383.53 | 1 |
| **Fructose** | 15.96 | 0.02 - 10 | y = 122359x - 15038 | 1 |
| **Fumaric acid** | 21.34 | 0.02 - 2.5 | y = 1E+06x - 37658 | 1 |
| **Galactose** | 15.41 | 0.02 - 20 | y = 98131x + 2788.4 | 1 |
| **Gluconic acid** | 14.48 | 0.08 - 10 | y = 251095x - 19919 | 1 |
| **Glucose** | 14.74 | 0.01 - 20 | y = 114349x - 1628.9 | 1 |
| **Glycerol** | 21.10 | 0.08 - 20 | y = 85957x + 12783 | 1 |
| **Glycolic acid** | 19.30 | 0.001 - 10 | y = 2E+06x + 263644 | 0.99 |
| **Glyoxylic acid** | 15.20 | 0.001 - 2.5 | y = 6E+06x + 106118 | 1 |
| **Inositol** | 15.27 | 0.02 - 20 | y = 134266x - 6021.2 | 1 |
| **Iso/Citric acid** | 13.29 | 0.02 - 20 | y = 497812x + 116819 | 1 |
| **Itaconic acid** | 20.27 | 0.0003 - 0.625 | y = 2E+07x + 191481 | 0.99 |
| **Lactose** | 12.86 | 0.02 - 10 | y = 110661x - 14488 | 1 |
| **Malic acid** | 15.35 | 0.08 - 10 | y = 530774x - 57569 | 1 |
| **Maltose** | 12.60 | 0.01 - 20 | y = 155198x - 3035.6 | 1 |
| **Mannitol** | 16.32 | 0.01 - 5 | y = 125447x - 1756.8 | 1 |
| **Mannose** | 15.50 | 0.01 - 20 | y = 123858x - 1377.8 | 0.99 |
| **Propionic acid** | 27.82 | 0.08 - 20 | y = 271649x - 69225 | 0.99 |
| **Quinic acid** | 15.97 | 0.02 - 20 | y = 273469x - 6796.8 | 1 |
| **Rhamnose** | 16.32 | 0.02 - 20 | y = 103262x + 1713.4 | 1 |
| **Ribose** | 17.41 | 0.02 - 20 | y = 94413x - 1881.9 | 0.99 |
| **Sorbitol** | 16.50 | 0.02 - 20 | y = 114021x - 7956.8 | 1 |
| **Sorbose** | 15.02 | 0.01 - 20 | y = 115937x + 104.75 | 0.98 |
| **Succinic acid** | 18.89 | 0.01 - 2.5 | y = 1E+07x + 272309 | 0.99 |

| | | | | |
|---|---|---|---|---|
| **Sucrose** | 12.66 | 0.02 - 20 | y = 57830x - 4731.2 | 1 |
| **Tartaric acid** | 13.88 | 0.01 - 20 | y = 724911x + 83746 | 0.99 |
| **Trehalose** | 12.55 | 0.005 - 20 | y = 207201x + 2227 | 1 |
| **Xylitol** | 17.76 | 0.01 - 20 | y = 135076x - 2438.7 | 1 |
| **Xylose** | 15.59 | 0.02 - 20 | y = 88764x - 3042.3 | 1 |

**Table S3:** Multiple Reaction Monitoring Library for MS/MS

| Compound ID | Ret. Time | Adduct ion | Polarity | Quantification ion | | | Reference ion 1 | | | Reference ion 2 | | | Quantification | | |
|---|---|---|---|---|---|---|---|---|---|---|---|---|---|---|---|
| | | | | Precursor ion | Product ion | CE (V) | Precursor ion | Product ion | CE (V) | Precursor ion | Product ion | CE (V) | Range (g/L) | Equation | $R^2$ |
| 2-ketogluconic acid | 12.49 | [M-H]- | (-) | 193 | 103 | 12 | 193 | 59 | 21 | 193 | 89 | 13 | 0.005 - 1.25 | y = 8E+06x - 74634 | 1 |
| Acetic acid | 23.93 | [M+H]+ | (+) | 61 | 43 | -10 | 61 | 41 | -11 | 61 | 44 | -18 | 0.005 - 20 | y = 779089x - 6696.4 | 1 |
| Aconic acid | 18.80 | [M-H]- | (-) | 173 | 111 | 10 | 173 | 129 | 11 | 173 | 111 | 10 | 0.04 - 5 | y = 87183x + 18055 | 0.99 |
| Adipic acid | 25.36 | [M-H]- | (-) | 145 | 101 | 14 | 145 | 83 | 14 | 145 | 81 | 21 | 0.02 - 5 | y = 2E+06x - 260622 | 1 |
| Alpha-ketoglutaric acid | 12.97 | [M+H]+ | (+) | 147 | 129 | -12 | 147 | 97 | -15 | 147 | 56 | -26 | 0.02 - 5 | y = 1E+06x + 24945 | 1 |
| Arabinose | 16.94 | [M+HCOO-]- | (-) | 195 | 89 | 12 | 195 | 149 | 10 | 195 | 59 | 22 | 0.01 - 5 | y = 238547x + 19731 | 0.99 |
| BDO | 32.84 | [M+H]+ | (+) | 91 | 43 | -17 | 91 | 55 | -12 | 91 | 73 | -9 | 0.01 - 2.5 | y = 4E+07x + 882321 | 1 |
| Cellobiose | 12.57 | [M+HCOO-]- | (-) | 387 | 341 | 8 | 387 | 161 | 12 | 387 | 101 | 22 | 0.01 - 1.25 | y = 159520x - 8273 | 1 |
| EG | 25.03 | [M+H]+ | (+) | 63 | 45 | -10 | 63 | 43 | -23 | 63 | 27 | -24 | 0.01 - 1.25 | y = 4E+06x + 156809 | 0.99 |
| Erythritol | 18.61 | [M+H]+ | (+) | 123 | 69 | -12 | 123 | 105 | -10 | 123 | 87 | -10 | 0.001 - 10 | y = 1E+07x - 70125 | 1 |
| Fructose | 15.96 | [M-H]- | (-) | 179 | 89 | 8 | 179 | 59 | 17 | 179 | 135 | 14 | 0.08 - 5 | y = 15645x + 5972.6 | 0.99 |
| Fumaric acid | 21.34 | [M-H]- | (-) | 115 | 71 | 10 | 115 | 98 | 22 | | | | 0.02 - 2.5 | y = 194879x - 12341 | 0.99 |
| Galactose | 15.41 | [M-H]- | (-) | 179 | 89 | 8 | 179 | 135 | 14 | 179 | 59 | 18 | 0.02 - 1.25 | y = 1541.2x + 2213.1 | 1 |
| Gluconic acid | 14.48 | [M-H]- | (-) | 195 | 129 | 13 | 195 | 75 | 17 | 195 | 99 | 14 | 0.04 - 10 | y = 686420x - 95583 | 1 |
| Glucose | 14.74 | [M+HCOO-]- | (-) | 225 | 179 | 8 | 225 | 89 | 13 | 225 | 59 | 22 | 0.01 - 2.5 | y = 473953x + 23292 | 0.99 |
| Glycerol | 21.10 | [M+H]+ | (+) | 93 | 57 | -11 | 93 | 75 | -9 | 93 | 45 | -14 | 0.001 - 10 | y = 6E+06x + 121362 | 1 |
| Glycolic acid | 19.30 | [M+HCOO-]- | (-) | 121 | 75 | 9 | 121 | 47 | 13 | 121 | 45 | 14 | 0.01 - 2.5 | y = 49705x + 3883.6 | 0.99 |
| Glyoxylic acid | 15.20 | [M-H]- | (-) | 73 | 45 | 10 | 73 | 29 | 11 | 73 | 29 | 11 | 0.08 - 2.5 | y = 3727.5x + 2955.4 | 0.98 |
| Inositol | 15.27 | [M+HCOO-]- | (-) | 225 | 179 | 11 | 225 | 45 | 23 | 225 | 161 | 18 | 0.04 - 5 | y = 61230x + 34912 | 0.99 |
| Iso/Citric acid | 13.29 | [M-H]- | (-) | 191 | 111 | 12 | 191 | 87 | 17 | 191 | 85 | 16 | 0.01 - 5 | y = 9E+06x - 323210 | 1 |
| Itaconic acid | 20.27 | [M+H]+ | (+) | 131 | 113 | -12 | 131 | 85 | -14 | 131 | 57 | -19 | 0.001 - 2.5 | y = 6E+06x + 63237 | 1 |
| Lactose | 12.86 | [M+HCOO-]- | (-) | 387 | 161 | 12 | 387 | 341 | 8 | 387 | 179 | 15 | 0.02 - 2.5 | y = 1E+06x - 47967 | 1 |
| Malic acid | 15.35 | [M-H]- | (-) | 133 | 115 | 15 | 133 | 71 | 15 | 133 | 73 | 16 | 0.01 - 10 | y = 1E+06x - 29988 | 1 |
| Maltose | 12.60 | [M+HCOO-]- | (-) | 387 | 161 | 12 | 387 | 341 | 8 | 387 | 179 | 15 | 0.16 - 20 | y = 18807x + 74334 | 0.99 |

| | | | | | | | | | | | | | | | |
|---|---|---|---|---|---|---|---|---|---|---|---|---|---|---|---|
| Mannitol | 16.32 | [M-H]- | (-) | 181 | 89 | 15 | 181 | 101 | 14 | 181 | 71 | 21 | 0.01 - 1.25 | y = 598007x - 6898.2 | 0.99 |
| Mannose | 15.50 | [M+HCOO-]- | (-) | 225 | 179 | 8 | 225 | 119 | 12 | 225 | 59 | 23 | 0.01 - 1.25 | y = 2E+06x + 151810 | 0.98 |
| Propionic acid | 27.82 | [M+H]+ | (+) | 75 | 57 | -13 | 75 | 29 | -16 | 75 | 27 | -24 | 0.005 - 20 | y = 6E+06x - 362268 | 1 |
| Quinic acid | 15.97 | [M-H]- | (-) | 191 | 85 | 21 | 191 | 93 | 22 | 191 | 127 | 18 | 0.01 - 10 | y = 710721x + 89781 | 0.99 |
| Rhamnose | 16.32 | [M-H]- | (-) | 163 | 59 | 13 | 163 | 103 | 8 | 163 | 89 | 6 | 0.16 - 2.5 | y = 20504x + 8738.4 | 0.98 |
| Ribose | 17.41 | [M+HCOO-]- | (-) | 195 | 149 | 10 | 195 | 89 | 12 | 195 | 45 | 23 | 0.02 - 10 | y = 211084x + 28765 | 0.99 |
| Sorbitol | 16.50 | [M-H]- | (-) | 181 | 89 | 14 | 181 | 101 | 15 | 181 | 71 | 21 | 0.312 - 20 | y = 129173x + 187056 | 0.99 |
| Sorbose | 15.02 | [M-H]- | (-) | 179 | 89 | 9 | 179 | 59 | 17 | 179 | 71 | 16 | 0.08 - 10 | y = 107485x + 63268 | 0.98 |
| Succinic acid | 18.89 | [M-H]- | (-) | 117 | 100 | 23 | 117 | 73 | 16 | 117 | 99 | 14 | 0.04 - 2.5 | y = 114364x - 93867 | 0.99 |
| Sucrose | 12.66 | [M+Na]+ | (+) | 365 | 203 | -23 | 365 | 185 | -21 | 365 | 23 | -40 | 0.001 - 1.25 | y = 3E+07x + 525075 | 1 |
| Tartaric acid | 13.88 | [M-H]- | (-) | 149 | 87 | 13 | 149 | 73 | 16 | 149 | 103 | 13 | 0.02 - 20 | y = 795107x + 43206 | 1 |
| Trehalose | 12.55 | [M+HCOO-]- | (-) | 387 | 341 | 13 | 387 | 179 | 19 | 387 | 89 | 26 | 0.001 - 0.312 | y = 9E+06x + 85156 | 0.98 |
| Xylitol | 17.76 | [M+H]+ | (+) | 153 | 107 | -13 | 153 | 79 | -16 | 153 | 61 | -30 | 0.005 - 1.25 | y = 1E+06x + 67560 | 0.98 |
| Xylose | 15.59 | [M-H]- | (-) | 195 | 89 | 12 | 195 | 149 | 10 | 195 | 59 | 21 | 0.04 - 20 | y = 146820x + 36513 | 0.99 |
